# Lysosomal reduced thiols are essential for mouse embryonic development

**DOI:** 10.1101/2024.12.26.630348

**Authors:** Charles H. Adelmann, Avanthika Venkatachalam, Lingjuan Huang, Michelle Liu, Sharon Germana, David M. Sabatini, David E. Fisher

## Abstract

While it has been appreciated for decades that lysosomes can import cysteine, its for organismal physiology is unclear. Recently, the MFSD12 transmembrane protein was shown to be necessary to import cysteine into lysosomes (and melanosomes), enabling the study of these processes using genetic tools. Here, we find that mice lacking *Mfsd12* die between embryonic days 10.5-12.5, suggesting that MFSD12 is essential for organogenesis. Within lysosomes, it is well known that a significant fraction of the cysteine is converted into cystine (oxidized cysteine), which accumulates and can be released into the cytosol via the CTNS (cystinosin) transporter. However, in contrast to *Mfsd12*, loss of *Ctns* results in live animals, indicating that it is not an essential gene. This suggests that the essential function of MFSD12 is not to enable cystine storage for use in the cytosol but rather to supply reduced cysteine to the lysosomal lumen itself. Consistent with this idea, the treatment of breeding heterozygous mice with cysteamine, a lysosome-penetrant thiol, rescued the development of *Mfsd12* knockout mice and resulted in live births. Our work implicates lysosomal thiol import as an essential metabolic pathway and provides tools for deciphering its complex genetic and metabolic interactions.

## Introduction

Proteins, nucleic acids, carbohydrates, and lipids are trafficked to the lysosome and broken down into their constitutive parts for reuse (1, 2) by the over 60 enzymes that operate within the carefully controlled environment of its lumen. A key metabolic feature of the lysosome is its thiol redox status. While the lysosome is overall oxidizing compared to the cytosol, proteins destined for degradation often have disulfide bonds that must be reduced to facilitate proteolysis, and the cysteine cathepsin proteases must maintain a reduced cysteine in their active sites to be functional (3–7).

Glutathione, the dominant reductive agent in the cytosol, does not appear to be transported by lysosomes (5, 6). Instead, lysosomes utilize cysteine for thiol reduction, importing it from the cytosol in a reduced form and exporting it as cystine after its oxidation (8–10). Two separate transport systems control lysosomal cysteine influx and cystine efflux and establish a pathway that has been well described in cell-free systems(9, 11). However, until recently the component(s) that comprise the cysteine import system were unknown, so it was not possible to genetically manipulate cysteine import to evaluate its significance *in vivo*.

While studying the human pigmentation gene *MFSD12*, we recently discovered that MFSD12, which is a 12 transmembrane domain protein, is not only required to import cysteine into melanosomes but also into the lysosomes of unpigmented cells (10). It is still unknown if the MFSD12 protein can directly transport cysteine when reconstituted into liposomes, but its function in cysteine transport appears quite central. Isolated lysosomes from *MFDS12* knockout cells have defective cysteine uptake, and relocalization of the MFSD12 protein to the plasma membrane is sufficient to introduce a new cell surface cysteine transport activity (10, 12). These findings establish MFSD12 as a promising genetic target to study the organismal importance of lysosomal cysteine.

Insights into the potential *in vivo* significance of lysosomal cysteine transport come from studies of CTNS (cystinosin), the well-established lysosomal cystine exporter (11, 13). Mutations in the *CTNS* gene cause the inherited disease cystinosis, which manifests in renal Fanconi syndrome, failure to thrive, and the progressive destruction of additional organ systems, pathologies thought to result from the buildup of cystine (13, 14). Recent studies examining the role of CTNS in models of development and cancer suggest that lysosomal cystine buffers cytosolic cysteine during oxidative stress and can influence pro-growth signaling (15–20). However, because the clinical features of cystinosis are thought to be driven in large part by mechanical damage from cystine crystals (13), it is challenging to interpret the disease phenotypes as purely caused by a lack of cystine export. Here, by comparing *Mfsd12* and *Ctns* knockout mice and manipulating lysosomal thiol content with cysteamine, we reveal lysosomal thiols as essential for embryonic development.

## Results

### Generation of an *Mfsd12* knockout allele in mice

We generated an *Mfsd12* mutant allele using CRISPR-Cas9 and an sgRNA designed to target all known isoforms of *Mfsd12* (Fig. 1A). The resulting 2-bp insertion (c.360_361insGA or *Mfsd12*^*em1Hadel*^) disrupts the 4th of the 12 predicted transmembrane domains in MFSD12 before generating a stop codon 28 amino acids downstream (Fig. 1B, Uniprot Q3U481). Because we initially intended to examine potential pigmentation phenotypes when we planned the generation of these mice, we backcrossed them with congenic C57BL/6J-*A*^*w-J*^ mice and bred white-bellied Jackson agouti (*A*^*w-J*^, red-brown and black fur) in place of *nonagouti* (*a*, black fur) (21). Consistent with the action of nonsense-mediated decay, *Mfsd12* expression was decreased to ~60% of wild-type levels in kidney and spleen extracts from *Mfsd12*^*em1Hadel*^ heterozygous mice (Fig. 1C).

**Figure 1.**
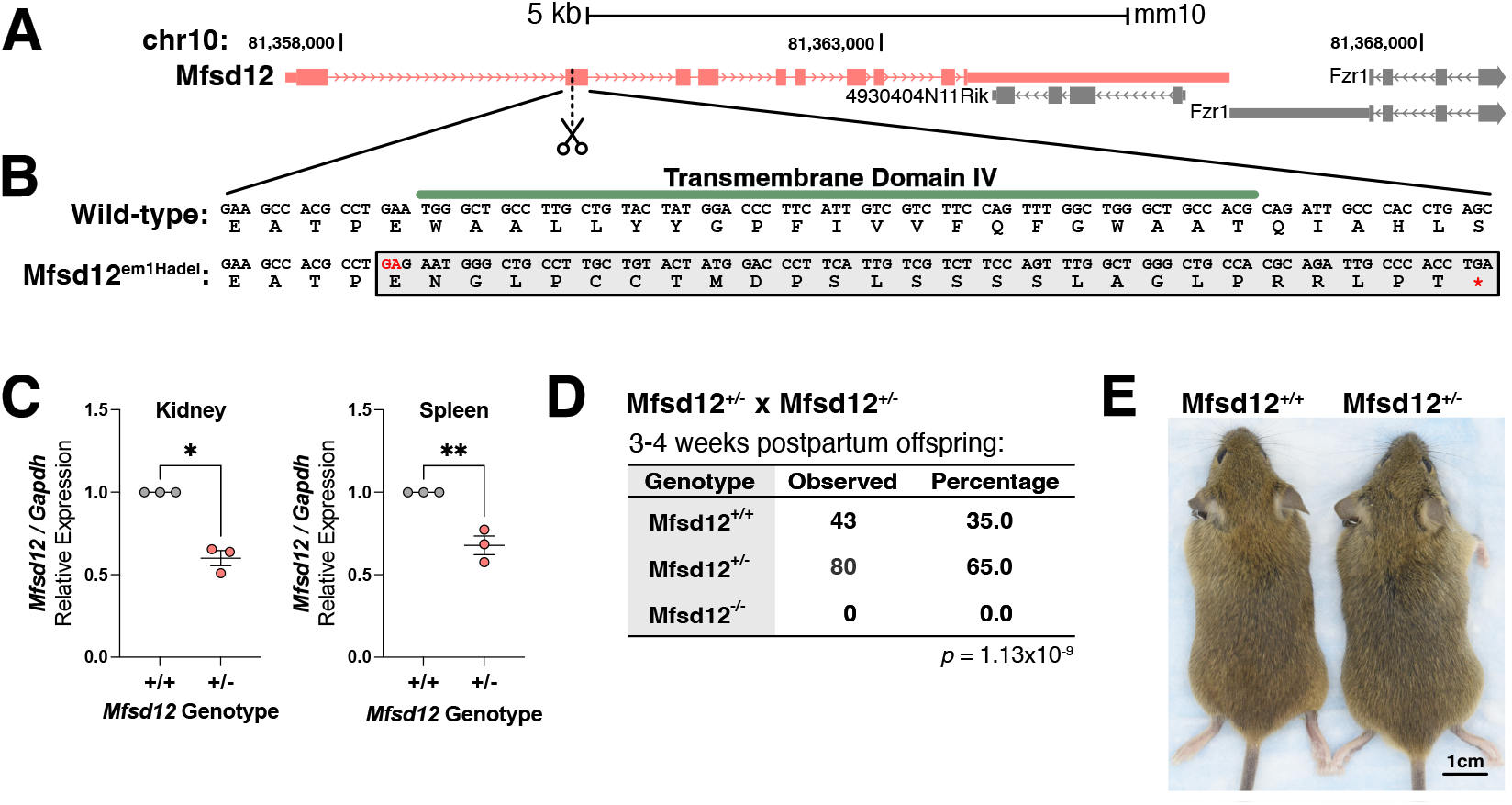
Loss of *Mfsd12* in mice results in completely penetrant lethality. **(A)** Diagram of murine *Mfsd12* gene structure and the location of the CRISPR-induced *Mfsd12* mutation. The diagram was accessed from the UCSC Genome Browser (mm10 genome build). **(B)** Nucleotide and amino acid resolution diagram of *Mfsd12*^*em1Hadel*^ mutation, showing the predicted amino acid sequence of the *Mfsd12* knockout allele. This includes an annotation of transmembrane domain IV from Uniprot Q3U481. Note the loss of hydrophobic amino acids (I, V, F etc.) **(C)** qRT-PCR analysis of *Mfsd12* expression in the kidney and spleen comparing *Mfsd12*^+/+^ and *Mfsd12*^+/−^ matched, male littermate pairs. Data are shown as mean ± SEM, with significance determined by unpaired, two-tailed Student’s t-test (\**p* < 0.05, \*\**p* < 0.01, n = 3 pairs of mice). **(D)** Offspring genotypes from 12 monogamous *Mfsd12*^+/−^ x *Mfsd12*^+/−^ crosses. The *p*-values are from a Chi-squared test for Mendelian ratios (n = 123 mice). **(E)** Representative photographs of 4-week-old *Mfsd12*^+/+^ and *Mfsd12*^+/−^ littermates show no obvious defects in body size or coat pigmentation.

We analyzed 12 separate monogamous crosses of *Mfsd12* heterozygous mice and despite weaning 123 mice at 3-4 weeks postpartum, we did not isolate a single *Mfsd12* knockout animal (Fig. 1D). Heterozygous animals appeared at a rate ~1.9-fold that of wild-type animals, suggesting loss of one copy of *Mfsd12* did not cause an intermediate viability phenotype. This is consistent with *Mfsd12* heterozygous mice having no detectable differences compared to wild-type counterparts in fur pigmentation or 4-week postpartum body weight (Fig. 1E and Fig. S1).

### *Mfsd12* knockout mice show development defects at 10.5-12.5 days post fertilization

To better understand why *Mfsd12* is essential, we sought to identify the stage at which *Mfsd12* knockout embryos cease to develop. Timed matings were set up with heterozygous *Mfsd12* mice, and embryos were isolated at defined time points before precise embryonic timing (“mE”) was further refined with development landmarks and computational analysis of limb morphology (22, 23). Knockout embryos isolated between embryonic day 10.5 and 12.5 were grossly abnormal (8 out of 9), with defects becoming most profound on the 12^th^ day of embryonic development (Fig. 2A-D). Early embryonic defects (mE10.5-11.5) were typically small size and reduced fetal blood (Fig. 2A), while later observed defects (mE12.0-13.0) included underdeveloped limbs and facial features (Fig. 2B). We interpreted these latter *Mfsd12* knockout embryos to be essentially dead, and consistent with this, very few *Mfsd12* knockout embryos were isolated after mE12.5, and none of these were normal (Fig. 2C and D).

**Figure 2.**
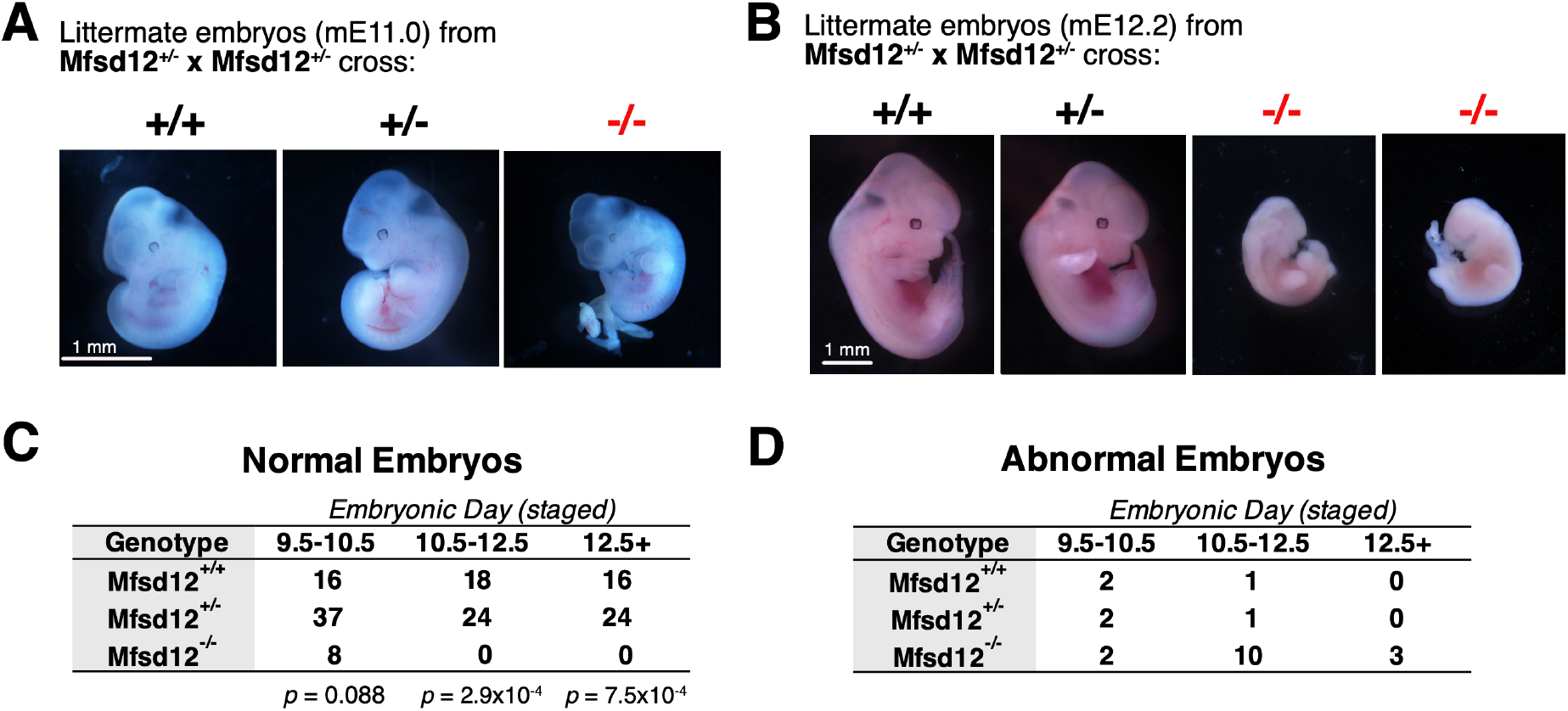
*Mfsd12* knockout induced embryonic lethality manifests 10.5-12.5 days after fertilization. **(A and B)** Photographs of littermate embryos staged at mE11.0 and mE12.2 illustrate the progressive defects in Mfsd12 knockout embryos. All *Mfsd12* knockout embryos shown were classified as “abnormal” based on size and morphology. Precise embryo staging was done by averaging limb staging data from eMOSS tool for “normal embryos” per litter (22, 23). **(C and D)** Tables showing the number of embryos isolated at indicated days post-fertilization and their genotypes. Embryos that were not visibly different from wild-type are included in Fig. 2C, while those that were deemed grossly abnormal based on morphology or small size are in Fig. 2D. The *p*-values are from a Chi-squared test for Mendelian ratios calculated for normal embryos (n = 61, 42, 40 embryos).

### The knockout of *Ctns* does not phenocopy that of *Mfsd12*

MFSD12 and CTNS act on two ends of a pathway, controlling cysteine and cystine movement through lysosomes (9, 10, 24). We hypothesized that by comparing loss-of-function phenotypes for both genes we should be able to understand if and how lysosomal cysteine is important for development. Several *Ctns* knockout alleles already exist, but we opted to generate a new one in parallel with *Mfsd12* to control for any confounding factors in genetic backgrounds and husbandry environments, as the former has been reported to affect both *Mfsd12* and *Ctns* phenotypes (14, 25).

Our *Ctns* targeting sgRNA introduced a 6-bp deletion compounded with a 1-bp insertion (Fig. 3A, c.329_335delinsT, *Ctns*^*em1Hadel*^). This frameshift mutation precedes the 1st transmembrane domain of the 7th transmembrane domain of the CTNS protein and results in a stop codon of 12 amino acids downstream (Fig. 3B, Uniprot 57757). Consistent with nonsense-mediated decay, we detected a decrease in *Ctns* levels to ~40-60% of wild-type levels in kidney and spleen tissues from *Ctns*^-/-^ mice (Fig. 3C).

**Figure 3.**
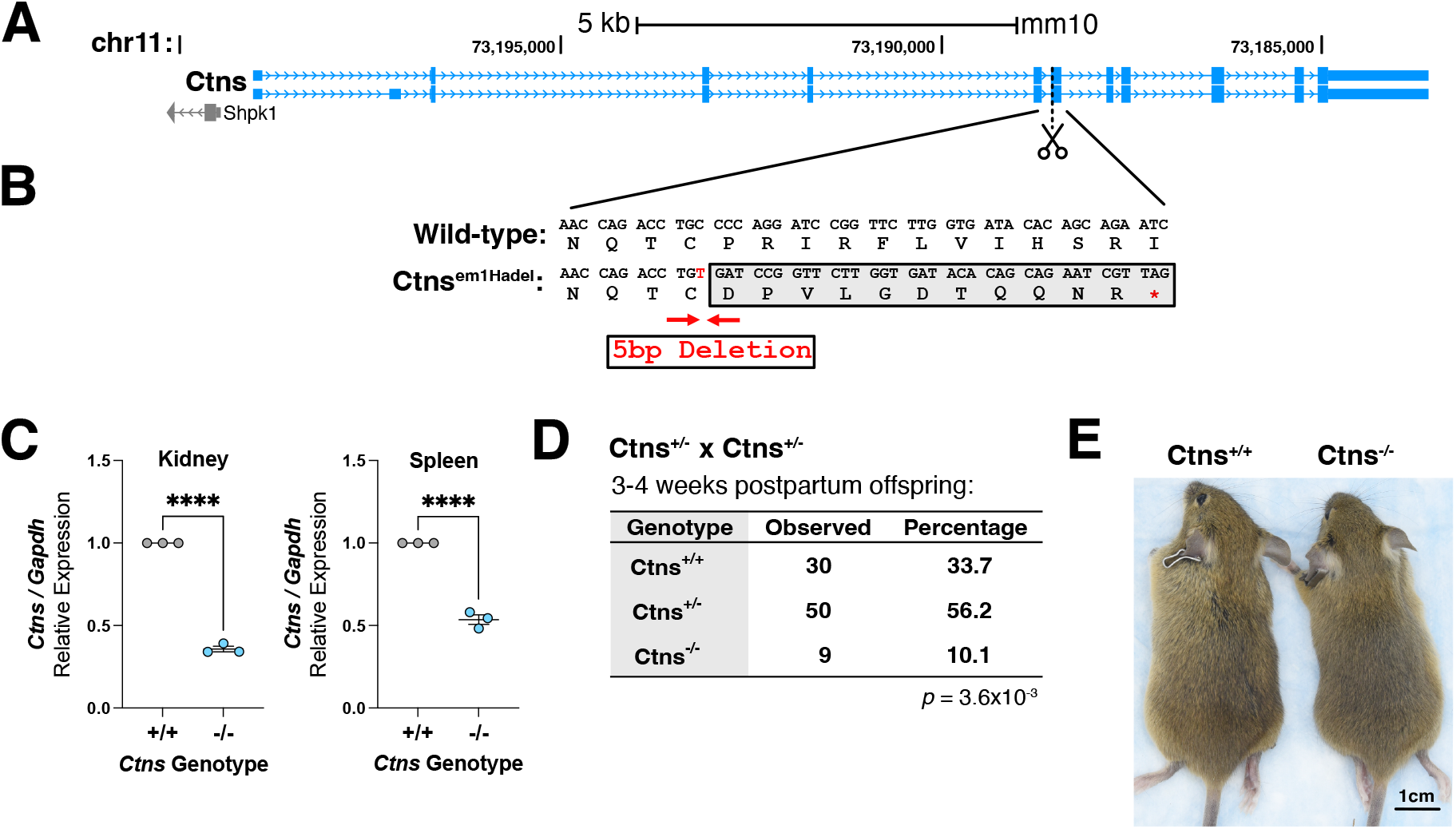
Knockout of *Ctns* does not phenocopy that of *Mfsd12* with respect to embryonic lethality. **(A)** Diagram of *Ctns* gene structure and the location of the CRISPR-induced *Ctns* mutation. The diagram was accessed from the UCSC Genome Browser (mm10 genome build) and is oriented in the antisense direction. **(B)** Nucleotide and amino acid resolution diagram of *Ctns*^*em1Hadel*^ mutation. **(C)** qRT-PCR analysis of *Ctns* expression comparing three *Ctns*^+/+^ and *Ctns*^-/-^ matched male littermate pairs. Data are shown as mean ± SEM, with significance determined by unpaired, two-tailed Student’s t-test (\*\*\*\**p* < 0.0001, n = 3 pairs of mice). **(D)** Number and frequency from 4 monogamous *Ctns*^+/−^ x *Ctns*^+/−^ crosses. The *p*-values are from a Chi-squared test for Mendelian ratios (n = 89 mice). **(E)** Photograph of 4-week-old *Ctns*^+/+^ versus *Ctns*^-/-^ littermates showing no obvious coat color differences.

After weaning 89 mice from 4 monogamous *Ctns*^+/−^ crosses, we isolated several live *Ctns* knockout mice 3-4 weeks postpartum. These knockout animals were isolated at a less-than-expected frequency (10% vs 25%; see Fig. 3D) and had reduced body weights (Fig. S2A). However, they were viable and could be used to breed further homozygous *Ctns* knockout mice (Fig. S2B). No pigment defects were observed in *Ctns* knockout animals, contrary to findings in other mouse models and in patients with cystinosis (Fig. 4D) (26).

**Figure 4.**
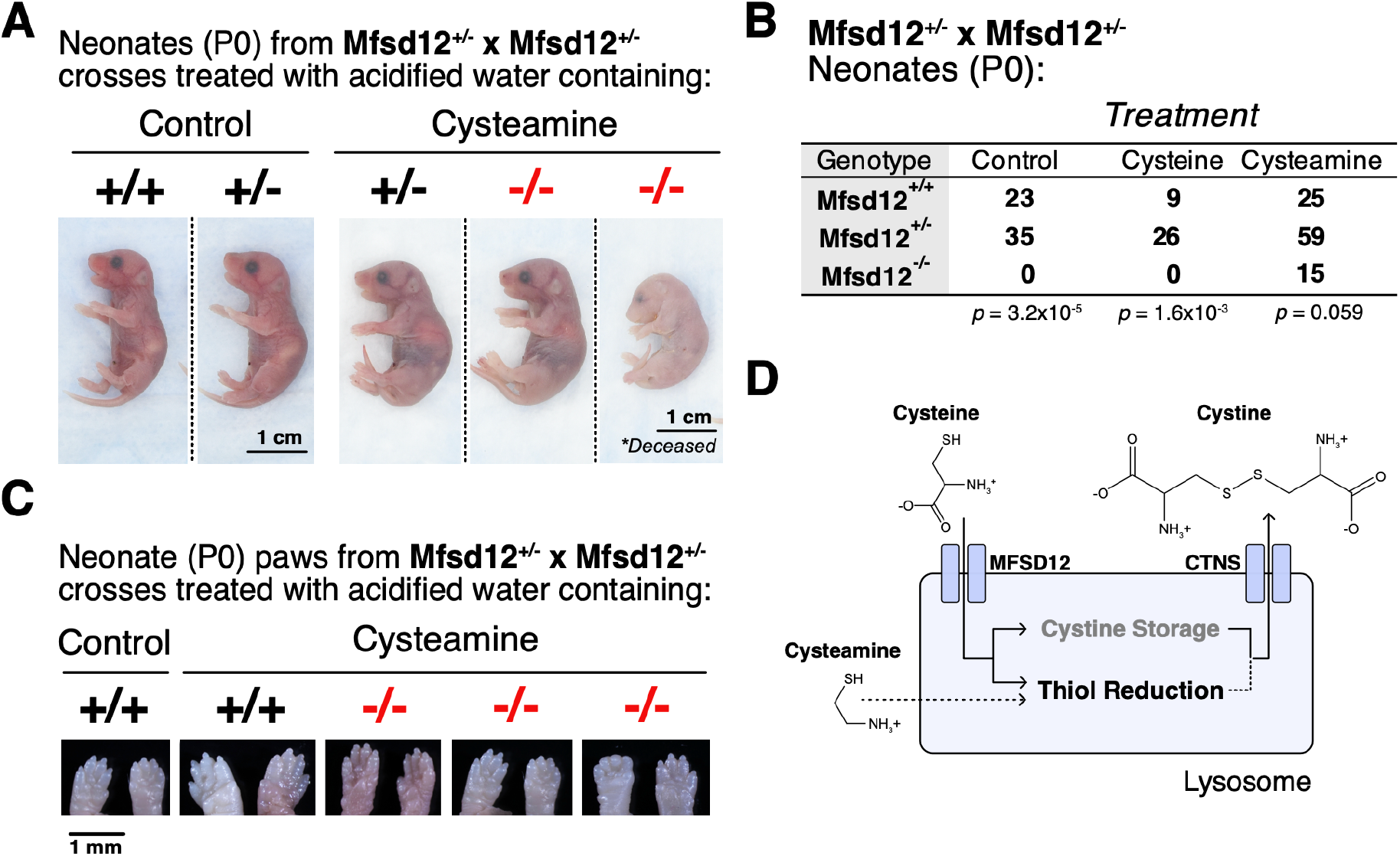
Lysosomal thiol supplementation with cysteamine rescues *Mfsd12* knockout animals to live birth. **(A)** Photograph of anesthetized neonates. Animals from the cysteamine-treated litter included an *Mfsd12*^-/-^ pup that was found deceased but is still typical of cysteamine rescue. For each treatment, littermates are shown. Photographs of control and cysteamine-treated mice were obtained on different days. **(B)** Genotype ratios of neonatal mice (both found decreased and alive) identified on the day they were born (P0) from mothers treated with control water (0.3% sucrose, pH 6.0), cysteine-supplemented water (8 mM cysteine, 0.3% sucrose, pH 6.0) or cysteamine-supplemented water (8 mM cysteamine, 0.3% sucrose, pH 6.0). The *p*-values are from a Chi-square test for Mendelian ratios (n = 58, 35, 99 neonates) **(C)** Neonate forepaws (ventral right and left paws) feature digits parallel to the midline and the presence of distal phalanges, features of development associated with progression past ~E16 (23). **(D)** Model of lysosomal cysteine and cystine metabolism in mouse embryonic development, highlighting the importance of luminal thiols like cysteine as reducing agents over cystine storage and release to the cytosol.

Efforts to breed *Mfsd12*^+/−^; *Ctns*^-/-^ mice to test for a direct genetic interaction between *Mfsd12* and *Ctns* were hindered by decreased offspring numbers (2.4 mice per litter measured over 5 litters from 3 breeding pairs) and the death of offspring before 3-weeks (10 out of 12 mice).

### Cysteamine treatment rescues the embryonic development of *Mfsd12* knockout mice to live birth

Our observation that the *Mfsd12* knockout does not phenocopy *Ctns* loss suggests that the essential function of MFSD12 in development is not to enable cystine storage for use in the cytosol via transport by CTNS. We reasoned instead that the *Mfsd12* knockout phenotype could be caused by the lack of reduced thiol import, a pathway shown *in vitro* to be important for degrading oxidized substrates in lysosomes (3–7).

To test this hypothesis, we treated *Mfsd12* heterozygous breeding pairs with cysteamine in their drinking water. Cysteamine is used in cystinosis therapy to convert cystine to cystine-cysteamine disulfides, which are effluxed from the lysosome through another transporter (27, 28). Cysteamine is highly bioavailable and well-documented to enter lysosomes in a transport-independent manner (13). Consistent with our embryonic studies, no newborn neonates (P0) were isolated from *Mfsd12*^*+/−*^ x *Mfsd12*^*+/−*^ breeding mice treated with a control drinking water (Fig. 4A and B, n = 58 mice). However, when we supplemented water with 8 mM cysteamine, knockout neonates were isolated at a frequency of 15% and at genotypic ratios that were overall consistent with Mendelian transmission (Fig. 4A and B, *p* = 0.059, Chi-square test for Mendelian transmission). As a control for adding free-thiol antioxidants to drinking water, we also treated breeding mice with 8 mM cysteine, which did not lead to the appearance of *Mfsd12* knockout neonates (Fig. 4A, n = 35 mice).

*Mfsd12* knockout neonates born to cysteamine-treated mothers had features indicating that they had advanced in development well past the stage at which the knockouts perished in the absence of cysteamine (Fig. 2A-B and Fig. 4A). These included well-developed eyes, skin wrinkling, and erupted distal phalanges that had converged to the midland of the forepaw (Fig. 4B-C and Fig. S3A). Skin development had also progressed to stages consistent with late embryogenesis, including cornification, panniculus carnosus formation, and hair follicle development (Fig. S3A)(29, 30). However, the rescue was not entirely complete as the *Mfsd12* knockout neonates weighed less than their wild-type and heterozygous counterparts from the cysteamine-treated crosses (Fig. S3B). Of the Mfsd12 knockout neonates, 6 out of 15 were found dead. While this was significantly elevated compared to the rate of dead neonates found with control treatment (3 out of 58, *p* = 2.0×10^-3^, Fisher’s exact test), it was not significantly different from the number of wild-type and heterozygous neonates found dead with cysteamine treatment (18 out of 84, *p* = 0.19, Fisher’s exact test). This suggested that cysteamine itself adversely affected embryonic development, birth, or perhaps early mother-offspring nursing interactions. Cysteamine-treated mice had no obvious change in hair color (Fig. S3C), suggesting that cysteamine, a known pigment-lightening agent (31, 32), might not have penetrated all tissues completely with our treatment regimen.

## Discussion

We find that in C57BL6/J-*A*^*w-J*^ mice, *Mfsd12* is a completely penetrant essential gene. Mice lacking functional *Mfsd12* progress through early embryonic development but die in the later stages associated with organogenesis. The essential role of MFSD12 is unlikely to be only to facilitate cystine storage or the flux of cysteine/cystine through the lysosome, as loss of *Ctns* results in live, viable mice. Instead, its essential role is likely to be in facilitating the use of cysteine as a lysosomal-reducing currency.

Consistent with this, treatment of pregnant mice with cysteamine rescued the embryonic embryonic death of *Mfsd12* knockout mice. Thus, lysosomal cysteine transport per se is necessary for animal embryonic development.

The defects we see in *Mfsd12* knockout mice are similar to the embryonic defects that have been previously reported in zebrafish embryos upon morpholino-mediated *mfsd12a* disruption (24), the partial embryonic lethality of intestine-specific *Mfsd12* knockout mice (12), and the embryonic lethality caused by whole-body loss of *Mfsd12* in C57BL/6J mice, which was reported without data (25). Interestingly, viable *Mfsd12* knockout mice have been reported in a mixed FVB/N x C57BL6/J background (25), suggesting that an outside variant can complement the role of MFSD12 in development (25). Biallelic frameshift mutations in *MFSD12* have also been reported in viable Shetland Ponies (33). Further analyses of the interactions between *Mfsd12* and other genetic loci might reveal new players in lysosomal cysteine metabolism.

The rescue by cysteamine of the development of *Mfsd12* knockout mice was profound but incomplete. Most of the Mfsd12 knockout neonates we isolated had reduced body weights and died during or soon after birth. Cysteamine undoubtedly oxidizes over time in the room-temperature drinking water used to deliver the compound to pregnant mice, and it has a short half-life in the organism, which is a known limitation of its use as a cystinosis treatment (13). Furthermore, the high rate of neonate death among wild-type and heterozygous neonates from cysteamine treatment suggests cysteamine-driven toxicities might synergize with the *Mfsd12* knockout developmental defects. The most interesting and not mutually exclusive implication is that a second mechanism operates downstream of *Mfsd12* but is not restored by cysteamine treatment. Recent work in cancer cell lines and flies suggests that lysosomal cystine stores can replenish cytosolic cysteine and buffer glutathione or acetyl-CoA under periods of oxidative stress (15–18, 20). The failure of the *Ctns* knockout to phenocopy that of *Mfsd12* suggests a cystine storage mechanism is not the primary reason *Mfsd12* is essential, but it does not rule out that it might play a contributing role.

The death of *Mfsd12* embryos after the cellular stages of development suggests that the essential role of *Mfsd12* is either cell type-specific or in the higher-order organization of tissues. The developmental defects we observed with germline *Mfsd12* knockout were extensive and did not immediately suggest the dysfunction of a single tissue system. Others have shown that *Vil1*-targeted deletion of *Mfsd12* interferes profoundly with the development of *Vil1* expressing tissues (intestines and eyes), but not in a manner that phenocopies our findings upon germline *Mfsd12* loss (12). These observations suggest that the *in vivo* defects elicited by *Mfsd12* loss might not have cell-type or tissue specificity but might result from a general feature of embryonic tissues, such as very rapid growth or a particular nutrient environment. We previously proposed that MFSD12 inhibition could be a therapy for cystinosis(10), and others working in zebrafish have noted the dose-limiting toxicity of *Mfsd12a* knockdown and the partial efficacy of its disruption in recovering renal function in *Ctns* mutant fish(24). Before further consideration of MFSD12 inhibition as a potential cystinosis therapy, the mechanism behind its essential function must be determined. For now, animal-based models like the one we present here provide a tractable system for untangling why *Mfsd12* is essential in tissues and organisms.

## Methods

### Mouse line generation and husbandry

Mice studies and procedures were approved by the Institutional Animal Care and Use Committees of the Whitehead Institute and Massachusetts General Hospital and were conducted strictly in accordance with the approved animal handling protocol. Animals were cared for according to the requirements of the National Research Council’s Guide for the Care and Use of Laboratory Animals.

C57BL/6J founder mice were generated with Cas9 and sgRNAs with the following targeting sequences:

Mfsd12: CGGGGAAGCCACGCCTGAAT

Ctns: TCACCAAGAACCGGATCCTG

After selecting frameshift alleles (Fig. 1A and Fig. 3A) for further study, mice were crossed to C57BL/6J-*A*^*w-J*^ (JAX#000051) to introduce the agouti coat color. Mice were genotyped using Sanger sequencing, next-generation amplicon sequencing (Quintara Biosciences), and a fluorescent probe-based qPCR assay implemented by TransnetYX interchangeably.

### *Mfsd12* and *Ctns* qRT-PCR

Mouse organs from sibling pairs were harvested, cut into pieces, and stored in an RNA preservative solution (saturated ammonium sulfate, 20 mM EDTA, 25 mM sodium citrate, pH 5.2) before storage at −80°C. Small pieces of tissue (~1×1 mm) were minced and placed in RLT Plus buffer (QIAgen, 1053393) supplemented with 140 mM 2-mercaptoethanol. Lysates were run through QIAshredder columns (QIAgen, 79656) before RNA was isolated via QIAgen RNAeasy Plus kit (QIAgen, 74134) according to the manufacturer’s instructions.

Gene expression was measured using the KAPA SYBR One-Step qRT-PCR kit (Sigma, SF1UKB) using the manufacturer’s protocol and the following PCR primers:

Mfsd12_F: GAACGAACCCAATGAGCACACC

Mfsd12_R: CTGAGACAGGTTCACAATGAGCC

Ctns_F: ATGGCTCCAGTTCCTCTTCTGC

Ctns_R: AATCGAGGAGCACACCGCCAAT

Gapdh_F: CTCCCACTCTTCCACCTTCG

Gapdh_R: GCCTCTCTTGCTCAGTGTCC

Target gene C_T_ values were determined automatically in QuantStudio Design Analysis 2 software (Applied Biosystems), normalized to *Gapdh*, and presented as a fold-change value determined via the ΔΔC_T_ method.

### Embryo harvest, staging, and characterization

Pregnant dams were sacrificed via CO_2_-based euthanasia at designated time points before harvest. Embryos were dissected in ice-cold PBS, and yolk-sacs or small tail sections were taken for genotyping before photo-documentation with a dissecting microscope. The precise embryonic day for each group of embryos was estimated with landmark features and with computational limb staging for embryos between mE10.25 and mE13.5 with eMOSS (https://limbstaging.embl.es)(22). For each litter of embryos, embryonic day values from “normal” embryos were averaged to estimate the reported age of the litter. Image scale bars were set by imaging a hemocytometer (Hausser Scientific) in parallel. Exposure conditions and brightness adjustments for each embryo image were not standardized.

### Supplementation of drinking water with cysteamine and cystine

200x cysteamine (Sigma Aldrich, M6500) and cysteine (Sigma Aldrich, 778451) solutions were made fresh in 60 mg/mL sucrose solutions and adjusted to pH 6.0 with sodium hydroxide before being diluted to 1X in vivarium-provided acidified water pH 6.0. Water was replaced every other day.

Cages were checked for neonates every morning. Neonates were euthanized, weighed, and fixed for 48 hours in a 10% formalin solution at room temperature. Pre-fixation, a 1-2 mm tail biopsy was taken for genotyping. Skin samples were taken from euthanized neonatal mice and embedded in paraffin using standard procedures. 5 μm sections were prepared from tissue blocks and stained with Hematoxylin Counterstain (Vector laboratories, H-3401) and Alcoholic Eosin Y Solution (Sigma, HT110132) for histologic analysis. Stained images were captured using a Hamamatsu Nanozoomer and analyzed using NDP.view2 software (Hamamatsu).

## Acknowledgments and Conflicts of Interest

We gratefully acknowledge the input of the entire Fisher lab, especially Judith R. Boozer and Jessica L. Flesher. Additionally, we are thankful to Raghu R. Chivukula for their input on the initial design of the mice and to Paul C. Rosen, Liron Bar-Peled, and Miranda Ravicz for critical discussion. We also thank the Whitehead Institute Genetically Engineered Models (GEM) Center for generating the mouse models in these experiments.

DEF acknowledges support to his laboratory from NIH grants P01 CA163222, R01 AR072304, and R01 AR043369, as well as funding from the Dr. Miriam and Sheldon G. Adelson Medical Research Foundation and the Melanoma Research Alliance. This work was also supported by the Leo Foundation to DMS (LF18057) and the Damon Runyon Cancer Research Fellowship to CHA (DG-2454-22).

CHA and DMS are listed on a patent (US20230103549A1) based on the work described here. DEF has a financial interest in Soltego, a company developing salt inducible kinase inhibitors for topical skin-darkening treatments that might be used for a broad set of human applications. The interests of DEF were reviewed and are managed by Massachusetts General Hospital and Partners HealthCare in accordance with their conflict-of-interest policies.

## Author Contributions

CHA designed the study and wrote the manuscript with input from DMS and DEF. SG helped design animal experiments, namely the breeding and timed mating setups. CHA and ML performed breeding experiments. CHA, AV, and LH performed molecular and histological characterizations.

