## Supplemental Figures for "Lysosomal reduced thiols are essential for mouse embryonic development"

**Supplemental Figures – Adelmann et al, 2024**

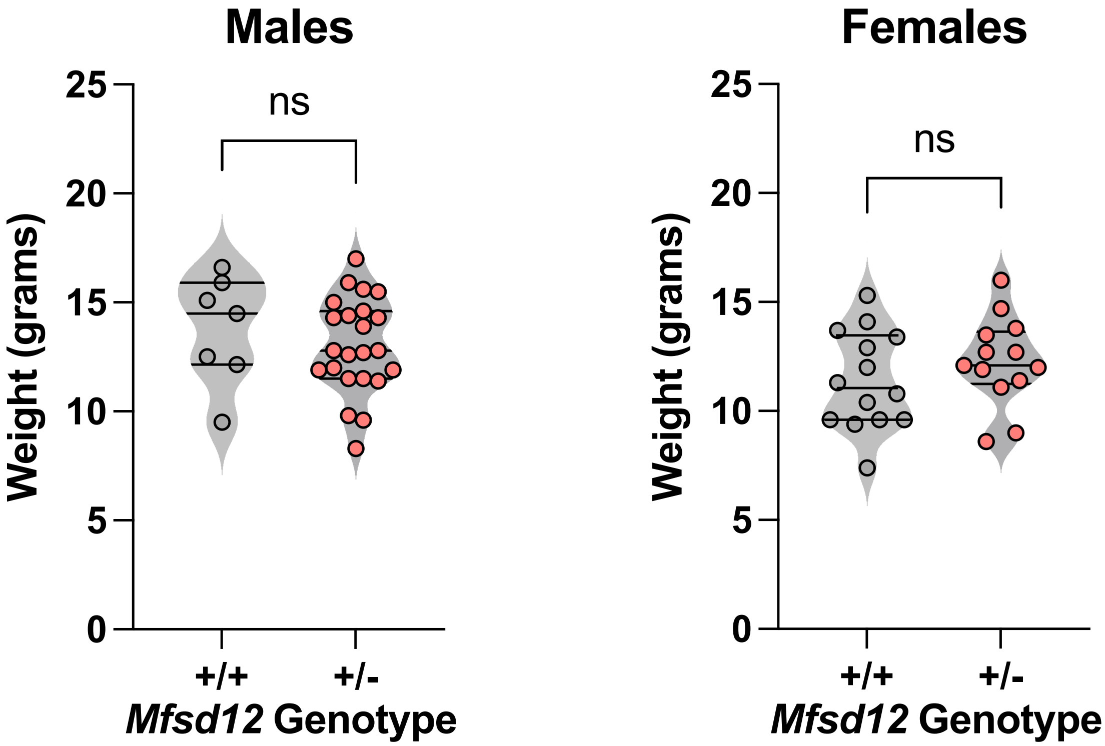


**Figure S1. *Mfsd12* heterozygous mice do not have defects in their birth weight.** Body weights of 4-week-old progeny from Mfsd12$^{+/-}$ x Mfsd12$^{+/-}$ crosses are shown, segregated by sex and genotype. Data are presented as median with interquartile range, with significance determined by unpaired, two-tailed Student's t-test ($^{ns}p > 0.05$, n = 7, 23, 14, and 13 mice for each genotype group, respectively).

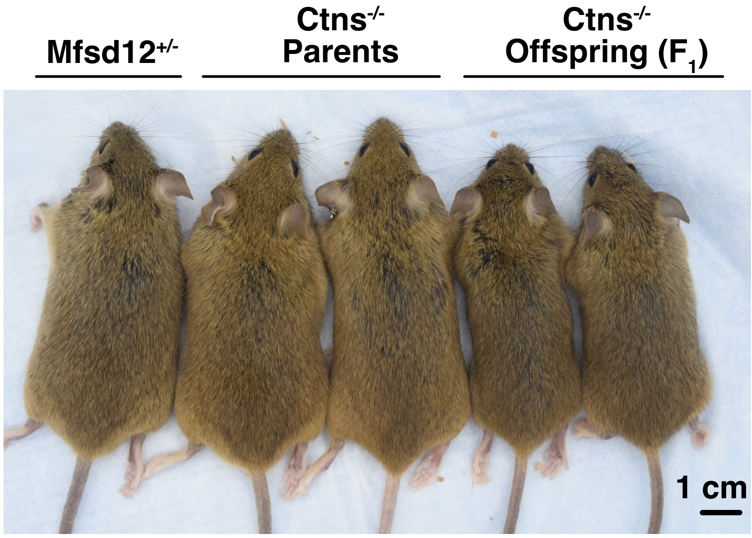


**Figure S2. *Ctns* knockout mice are viable and propagate to the next generation.** Photograph of *Mfsd12*$^{+/-}$ mouse (included as a control), *Ctns*$^{-/-}$ parents, and their direct progeny.

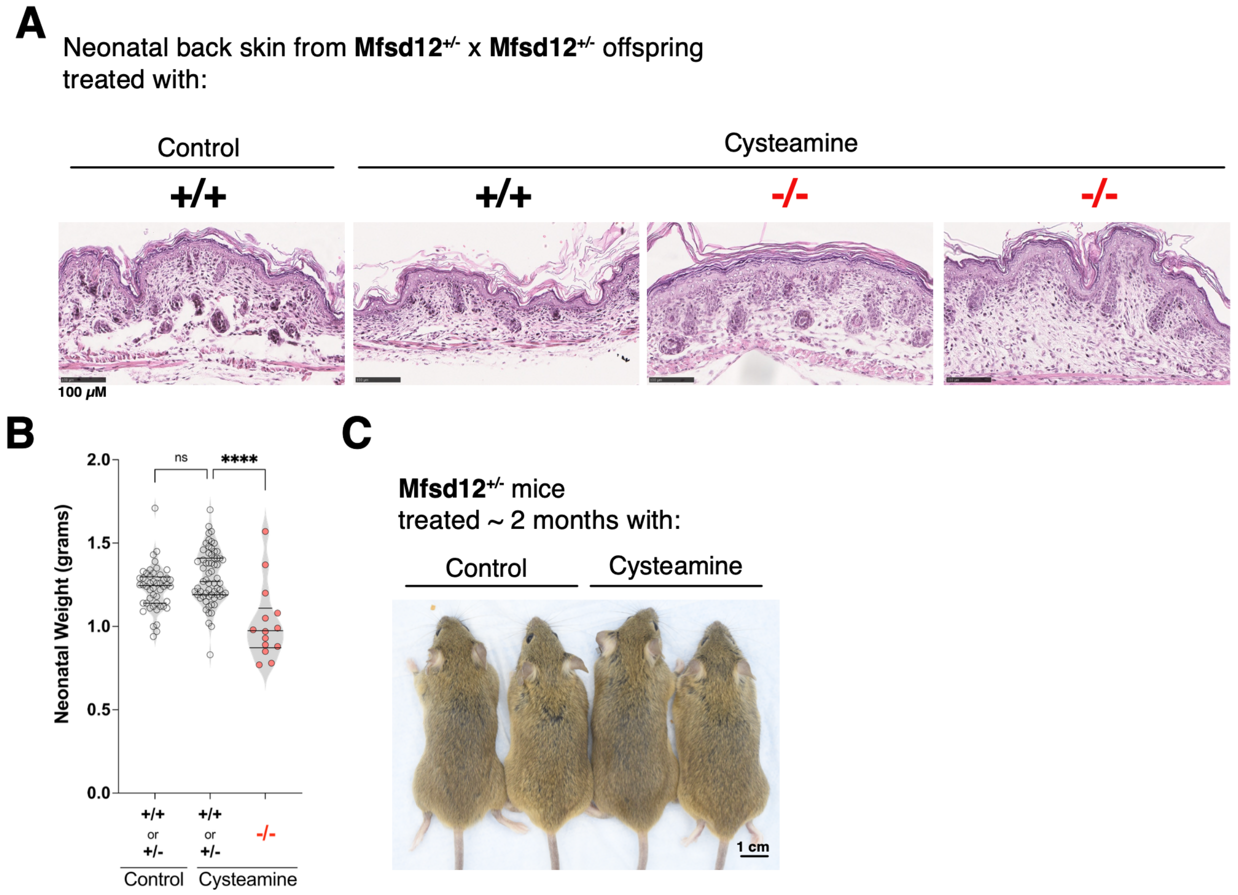


**Figure S3. Cysteamine rescues the development of *Mfsd12* knockout mice. (A)** Representative images of neonatal skin from control and cysteamine-treated animals, showing cornification, panniculus carnosus formation, and follicle development in cysteamine-rescued *Mfsd12* knockout skin. **(B)** Body weights of neonates from control and cysteamine-treated breeding mice, organized by genotype. Neonates with evidence of chewing or biting trauma were excluded from the analysis. Data are shown as median with interquartile range with statistical significance determined by one-way ANOVA (****$p < 0.0001$, n = 48, 63, and 15 neonates for each group, respectively). **(C)** Cysteamine-treated males from experimental breeding mice after ~2 months of cysteamine treatment.
